# Genome-wide analysis of Structural Variants in Parkinson’s Disease using Short-Read Sequencing data

**DOI:** 10.1101/2022.08.22.504867

**Authors:** Kimberley J. Billingsley, Jinhui Ding, Pilar Alvarez Jerez, Anastasia Illarionova, Francis P. Grenn, Mary B. Makarious, Anni Moore, Daniel Vitale, Xylena Reed, Dena Hernandez, Ali Torkamani, Mina Ryten, John Hardy, UK Brain Expression Consortium (UKBEC), Ruth Chia, Sonja W. Scholz, Bryan J. Traynor, Clifton L. Dalgard, Debra J. Ehrlich, Toshiko Tanaka, Luigi Ferrucci, Thomas.G. Beach, Geidy E. Serrano, John P. Quinn, Vivien J. Bubb, Ryan L Collins, Xuefang Zhao, Mark Walker, Emma Pierce-Hoffman, Harrison Brand, Michael Talkowski, Bradford Casey, Mark R Cookson, Androo Markham, Mike Nalls, Medhat Mahmoud, Fritz J Sedlazeck, Cornelis Blauwendraat, J. Raphael Gibbs, Andrew B. Singleton

## Abstract

Parkinson’s disease is a complex neurodegenerative disorder, affecting approximately one million individuals in the USA alone. A significant proportion of risk for Parkinson’s disease is driven by genetics. Despite this, the majority of the common genetic variation that contributes to disease risk is unknown, in-part because previous genetic studies have focussed solely on the contribution of single nucleotide variants. Structural variants represent a significant source of genetic variation in the human genome. However, because assay of this variability is challenging, structural variants have not been cataloged on a genome-wide scale, and their contribution to the risk of Parkinson’s disease remains unknown. In this study, we 1) leveraged the GATK-SV pipeline to detect and genotype structural variants in 7,772 short-read sequencing data and 2) generated a subset of matched whole-genome Oxford Nanopore Technologies long-read sequencing data from the PPMI cohort to allow for comprehensive structural variant confirmation. We detected, genotyped, and tested 3,154 “high-confidence” common structural variant loci, representing over 412 million nucleotides of non-reference genetic variation. Using the long-read sequencing data, we validated three structural variants that may drive the association signals at known Parkinson’s disease risk loci, including a 2kb intronic deletion within the gene *LRRN4*. Further, we confirm that the majority of structural variants in the human genome cannot be detected using short-read sequencing alone, encompassing on average around 4 million nucleotides of inaccessible sequence per genome. Therefore, although these data provide the most comprehensive survey of the contribution of structural variants to the genetic risk of Parkinson’s disease to date, this study highlights the need for large-scale long-read datasets to fully elucidate the role of structural variants in Parkinson’s disease.

## Introduction

Although the exact cause of Parkinson’s Disease is unclear, there is substantial evidence that genetic factors contribute to the risk of developing the disease. To note, the most recent genome-wide association study (GWAS) which included around 40k cases, 20k proxy-cases and 1.4 million controls, identified 90 independent risk signals across 78 regions of the genome ^1^. Despite these large-scale efforts, there remains two major gaps in our understanding of the genetics of Parkinson’s Disease risk; 1) we have a limited understanding of what variants and genes are driving the signal at the known risk loci and 2) if taken together, these loci explain 16-30% of the heritable component of the disease, meaning that the majority of the common genetic variation that contributes to disease risk is yet to be discovered ^1^. This is often termed the “missing-heritability” of Parkinson’s Disease.

Previous large-scale genetic studies have focused solely on single-nucleotide variants (SNVs), which represent only a fraction of the genetic variation in the human genome. Structural variants (SVs), here defined as any DNA rearrangements involving at least fifty nucleotides, represent over ten times more genetic variation than SNVs^2^. Using array-based and short read sequencing (SRS) datasets, SVs are more difficult to identify and accurately genotype than SNVs, the detection of SV being confounded by common sequencing and alignment artifacts. As a result of this challenge, SV have been largely understudied. The development of high-throughput sequencing technologies that have allowed for widespread use of whole genome sequencing (WGS) in population genetics, along with advances in scaling SV detection algorithms, have combined to allow an initial investigation of SVs for large SRS WGS cohorts.

SVs are typically classified as deletions, duplications, insertions, inversions, and translocations (abbreviated to DEL, DUP, INS, INV, CTX respectively) describing different combinations of gains, losses, or rearrangements of DNA sequence ^3–5^. SVs also include; copy number variants (CNV), mobile element insertions (MEI), such as *Alu*, LINE-1 and SINE-VNTR-*Alu* (SVA) elements that can still insert themselves across the human genome and complex SVs (CPX) that consist of multiple combinations of these described events. These genetic variants can have a substantial phenotypic impact, disrupting gene function and regulation or modifying gene dosage. Further, recent studies have shown that SVs drive functional changes across populations and cell and tissue types ^6–8^.

Although the role of SVs is yet to be comprehensively assessed in the context of risk of apparently sporadic Parkinson’s Disease on a genome-wide scale, early studies using targeted approaches discovered several SVs that are causative of monogenic forms of Parkinson’s Disease and Parkinsonism. Examples of this include partial deletions in the gene *PARK2* identified as causative for autosomal recessive Parkinson’s Disease^9^ and then the major discovery of a CNV of the entire genomic region encompassing the gene *SNCA* that was shown to be causative of a form of autosomal dominant Parkinson’s Disease in a large family^10^. Since then, causative CNVs have been reported in other familiar Parkinson’s Disease genes, such as the genes *PINK1*^11^ and *PARK7*^12^. In addition, a MEI SVA insertion within intron 32 of the gene *TAF1* is causative of X-Linked Dystonia Parkinsonism^13–15^ and variation within the SVAs hexanucleotide repeat domain is associated with age of disease onset and influences transcriptional activity of *TAF1*^16^.

In this study, we focussed on common bi-allelic SVs and performed the first genome-wide characterization of SVs in Parkinson’s Disease to date. In 7,772 SRS genomes, we detected a total of 227,357 SVs and validated several SVs associated with Parkinson’s Disease risk at known loci. Finally, we highlight that the majority of SVs in the genome cannot be captured by SRS alone.

## Materials and methods

### Samples and quality control

#### Parkinson’s Disease and control cohort description

The following ten cohorts were used in this study; Biofind (https://biofind.loni.usc.edu/), Harvard Biomarkers Study (HBS) (https://amp-pd.org/unified-cohorts/hbs), North American Brain Expression Consortium (NABEC)^17^, Laboratory of Neurogenetics pathologically confirmed collection, the NINDS Parkinson’s Disease Biomarker Program (PDBP) (https://pdbp.ninds.nih.gov/), samples from the NIH Parkinson’s Disease Clinic (NIH PD CLINIC), the Parkinson’s Progression Markers Initiative (PPMI) (https://www.ppmi-info.org/), Wellderly (controls), and the United Kingdom Brain Expression Consortium (UKBEC). Clinical and demographic characteristics of the cohorts under study are shown in (Supplementary Table 1). Participants included Parkinson’s Disease cases clinically diagnosed by experienced neurologists. All Parkinson’s Disease cases met criteria defined by the UK Parkinson’s Disease Society Brain Bank^18^.

#### Whole genome sequencing

SRS data generation through AMP-PD has been reported in detail previously by Iwaki *et al^19^*, but in brief DNA sequencing was performed using two providers, Macrogen and Uniformed Services University of the Health Sciences (USUHS). For samples sequenced at Macrogen one microgram of each DNA sample was fragmented by Covaris System and further prepared according to the Illumina TruSeq DNA Sample preparation guide to obtain a final library of 300-400 bp average insert size. Libraries were multiplexed and sequenced on the Illumina HiSeq X platform. For samples sequenced by USUHS, DNA samples were processed using the Illumina TruSeq DNA PCRFree Sample Preparation kit, starting with one microgram input and resulting in an average insert size of 410 bp. USUHS sequenced DNA samples were run single-library, single lane on a HiSeq X flowcell (Illumina).

#### Sequence alignment and single nucleotide variant calling

Paired-end read sequences were processed in accordance with the pipeline standard developed by the Centers for Common Disease Genomics^20^. The GRCh38DH reference genome was used for alignment as specified in the standardized functional equivalence (FE) pipeline^21^. The Broad Institute’s implementation of this FE standardized pipeline, which incorporates the GATK (2016) Best Practices^22^, is publicly available and used for WGS processing. Single-nucleotide (SNV) and indels were called from the processed WGS data following the GATK (2016) Best Practices^22^ using the Broad Institute’s workflow for joint discovery and Variant Quality Score Recalibration (VQSR) ^23^. For quality control, each sample was checked using common methods for genotypes and sequence related metrics. Using Plink v1.9^24^ each sample’s genotype missingness rate (< 95%), heterozygosity rate (exceeding +/− 0.15 F-stat), and gender were checked. The King v2.1.3^25^ kinship tool was used to check for the presence of duplicate samples. Sequence and alignment related metrics generated by the Broad’s implementation of the FE standardized pipeline were inspected for potential quality problems. This included the sample’s mean sequence depth (< 30X) and contamination rate (> 2%), as reported by VerifyBamID^26^, and single nucleotide variant count as reported by Picard’s CollectVariantCallingMetrics (< 3 StDev) based on the sample’s genomic VCF (gVCF). Principal components (PCs) were created for each dataset using PLINK. For the PC calculation, variants were filtered for minor allele frequency (< 0.01), genotype missingness (< 0.05) and Hardy–Weinberg equilibrium (P <= 1E-6) and pruned (--indep-pairwaise 1000 10 0.02).

#### Structural Variant discovery with the GATK-SV pipeline

For SV discovery and downstream filtering, the Broad Institute GATK-SV pipeline was run in cohort mode https://github.com/broadinstitute/gatk-sv^27^. All computations were finished on the Google Cloud Platform (https://cloud.google.com).

#### Sample Quality Control

For each sample, 10 quality control features were measured, including median sequencing coverage in 100bp bins, dosage bias score δ, autosomal ploidy spread, Z-score of outlier 1Mb bins, chimera rate, pairwise alignment rate, read length, library contamination, ambiguous sex genotypes, and discordant inferred and reported sex. We filtered the dataset with these 10 measurements and excluded 164 samples (2.03%) for failing at least one criterion. We kept samples with non-canonical sex chromosome configurations in their batches and manually removed all raw SV calls on X/Y from their raw VCF files.

#### Sample batching

To facilitate SV discovery computing on Google Cloud Platform and control for potential batch effects and confounders, we followed the batch scheme designed by Collins *et al* to subdivide all 7,914 samples that passed QC into 20 batches with ~400 samples per batch^27^. All samples were sequenced with a PCR-free protocol: first, samples were assigned to a female group or a male group based upon their chrX ploidy estimates; second, each gender group was further split into 4 subgroups based upon their median 100bp binned coverage values; third, in each subgroup, samples were ranked using dosage bias score and further partitioned into bundles of ~100 samples, with the number of bundles of female samples being equal to those of male samples; finally, we merged every two batches of ~100 male samples with two batches of ~100 female samples that corresponded in coverage rankings and dosage bias scores. The ratio of female samples and male samples in each batch is around 1:1.19, balanced across all batches.

#### Structural Variant evidence collection

The SV evidence of individual samples collected in this step include: raw SV calls from three different structural variant (SV) algorithms Manta v1.4^28^, MELT v2.2.0 ^29^, and Wham v1.7 ^30^; CNV calls using cn.MOPS v1.20.1 ^31^ and GATK gCNV^32^ from depth evidence; and raw evidence binned read counts (RD), split reads (SR), discordant read pairs (PE), and SNP B-allele frequency (BAF). We then aggregated the SV calls from the three algorithms and the CNV calls of each sample and standardized the calls to meet specifications required for the SV discovery pipeline. The computations in this step were executed on Google Cloud Platform, and all evidence was collected using the GATK-SV pipeline, except the SV calls from Manta and MELT, which we collected separately. MELT Single was run with a custom “priors” vcf with parameter (-r) set to a read length of 150bp. The custom priors vcf contained MEI sites that were discovered when MELT was run with a subset of the PPMI samples.

#### Structural Variant discovery

The core SV discovery steps include: clustering SV calls across every batch; generating four variant matrices of each batch, corresponding to Manta, MELT, Wham, and depth-based callers (cn.MOPS and GATK gCNV), respectively; filtering low quality variants and outlier samples in each batch; combining filtered variants across batches; genotyping per batch samples across unfiltered variants combining 20 batches; re-genotyping probable variants across all batches; combining variants across 20 batches; resolving complex variants; and conducting a final VCF file clean up. For the GATK gCNV analysis, gCNV models were trained for each batch on 100 samples randomly selected from the batch. All steps were included in the GATK-SV pipeline, which we executed in cohort mode while strictly following the default parameters settings.

#### Structural Variant filtering

We ran all four downstream filtering steps included in the GATK-SV pipeline: minGQ filtering, FilterOutlierSamples, BatchEffect, and FilterCleanupQualRecalibration. Since the Parkinson’s Disease cohort lacks family structures, we ran minGQ filtering using the table pre-trained with 1000 genomes samples at the 1% FDR thresholds stored in the gatk-sv public resources bucket on Google Cloud Platform (https://gs://gatk-sv-resources-public/hg38/v0/sv-resources/ref-panel/1KG/v2/mingq/1KGP_2504_and_698_with_GIAB.1perc_fdr.PCRMINUS.minGQ.filter_lookup_table.txt). 142 outliers were removed in the FilterOutlierSamples step, executed the procedures of BatchEffect, and ran FilterCleanupQualRecalibration on the remaining cleaned 7772 samples. The final SV call set included 366,555 SV calls.

### Structural Variant association analyses

#### Sample Quality control

Initial sample inclusion criteria included; age at disease onset or last examination at 18 years of age or older, no genetically ascertained relation to other samples (proportional sharing at a maximum of 12.5%) at the cousin level or closer and majority European ancestry confirmed through principal-components determined by HapMap3. All individuals recruited as part of a biased and/or genetic cohort, such as *GBA* and *LRRK2* rare variant carriers within a specific effort of PPMI cohort, were also excluded.

After sample QC, a total of 2,585 Parkinson’s Disease cases and 2,779 neurologically healthy controls were included. Parkinson’s Disease cases ranged from 19 to 92 years of age of onset. Control subjects ranged from 19 to 110 years of age.

#### Variant Quality control

For the association analyses in order to assess the impact of high-quality variants, SVs with the filter label “PASS” were extracted from the final SV call set leaving a total of 227,357 biallelic automatal SVs.

#### Statistical analysis

We performed a SV Parkinson’s Disease GWAS (n= 2,585 cases, 2,779 controls) using logistic regression in PLINK(v2.0) with a minor allele frequency threshold of >1%. Principal components (PCs) were generated in PLINK(v1.9) for the common SNV datasets and common SV dataset separately. The step function in R MASS package was used to identify the minimum number of PCs required to correct for population substructure^24^ using both sets of PCs. Based on this analysis sex, age, and 18 PCs (SNV_PC1, SNV_PC2, SNV_PC4, SNV_PC5, SNV_PC6, SNV_PC7, SNV_PC8, SNV_PC9, SNV_PC10, SV_PC1, SV_PC2, SV_PC3, SV_PC4, SV_PC5, SV_PC6, SV_PC8, SV_PC9, SV_PC10) were incorporated in the model. Overall genomic inflation was minimal with a lambda estimate of 1.011 and a lambda scaled to 1,000 cases and 1,000 controls at 1.004. Multiple test correction was handled using standard Bonferroni correction in PLINKv1.9 under default settings.

#### Linkage Disequilibrium tagging for Structural Variants

Given that we do not understand what variant is driving the known Parkinson’s Disease association signals and that previous studies have shown that SVs drive known GWAS associations for other diseases, we next integrated the new SV dataset with the corresponding SNV data to identify if any SV tags any of the 90 Parkinson’s Disease risk SNVs. LD between SNVs and SVs was computed with the “--r2 inter-chr dprime” parameter in Plink v1.9^33^.

#### Extracting variants within Parkinson’s Disease genes

To identify variants within Parkinson’s Disease causal genes that were only present in Parkinson’s Disease cases, 21 genes were included whereby mutations have been recorded to cause Parkinson’s Disease. The following genes were included; *SNCA, PRKN, UCHL1, PARK7, LRRK2, PINK1, POLG, HTRA2, ATP13A2, FBX07, GIGYF2, GBA, PLA2G6, EIF4G1, VPS35, DNAJC6, SYNJ1, DNAJC13, TMEM230, VPS13C* and *LRP10*. Genomic coordinates were defined using Refseq. GATK-SV “PASS” variants were intersected with the 21 regions with bedtools ^34^ “–intersect”.

### Structural Variant Confirmation with Long-Read Sequencing

#### Nanopore Long-Read Sequencing from the PPMI blood samples

Matched ONT LRS was generated for 8 PPMI samples from blood to *in-silico* validate the SRS discovered SV of interest. The samples were processed and sequenced using a protocol optimized for population scale long-read sequencing from frozen human blood (https://dx.doi.org/10.17504/protocols.io.kxygxzmmov8j/v2). In brief, high molecular weight DNA was extracted from 1mL of blood using the Kingfisher APEX instrument and the Nanobind CBB Big DNA kit (HBK-CBB-001) with 4mm Nanobind disks from Circulomics. Next, the DNA went through a size selection step using the Circulomics Short Read Eliminator Kit (SS-100-101-01) to remove fragments up to 25kb. DNA was then sheared to a target size of 30 kb using a Megaruptor 3 instrument and the standard Diagenode Megaruptor 3 shearing kit. A library was prepared with the SQK-LSK 110 Ligation Sequencing Kit from ONT. The samples were quantified using a Qubit fluorometer and were loaded onto a PromethION R9.4.1 flow cell following ONT standard operating procedures and ran for a total of 72 hours on a PromethION device.

#### Long-read Sequencing data processing

Fast5 files containing raw signal data were obtained from sequencing performed using minKNOW v21.05.13 (Oxford Nanopore Technologies). All fast5 files were used to perform super accuracy basecalling on each sample with Guppy v6.0.1 (Oxford Nanopore Technologies) using the following command:

guppy_basecaller -i ${FAST5_PATH} -s ${OUT_PATH} -c dna_r9.4.1_450bps_sup_prom.cfg-x cuda:all:100% -r --read_batch_size 250000 -q 250000

Prior to mapping with winnowmap v2.03^35^, meryl was used to identify and count the top 0.02% most frequent 15-mers in the hg38 reference genome using the following commands:

meryl count k=15 output merylDB GRCh38.primary_assembly.genome.fa

meryl print greater-than distinct=0.9998 merylDB > repetitive_k15.txt

Fastq files that passed quality control filters in the super accuracy basecalling step were then mapped to the GRChg38 reference genome using the following winnowmap command:

winnowmap -W repetitive_k15.txt -ax map-ont GRCh38.primary_assembly.genome.fa sample_pass1.fastq.gz > sample_pass1.sam

The resulting sam files were sorted, converted to bams and indexed using samtools^36^ and were then merged, sorted and indexed with samtools to create one final bam file per sample. Chimera rate was calculated using the Liger2LiGer tool ^37^.

#### Long-read Sequencing data Structural Variant discovery and comparison

Sniffles2 ^38^ v2.0.3 was run using default parameters to discover and genotype SVs in the matched LRS data. To improve SV calling in repetitive regions, the “–tandem-repeats’’ option was used. To filter out possible false positive SV calls Survivor^39^ v1.0.7 was used with the” –filter” option to remove SV below 50bp.

Additionally, to identify if there was evidence of the GATK-SV predicted SV in the matched LRS, we extracted only predicted non-reference SV carriers with the Samtools^40^ v1.3 “--min-ac=1” parameter. Next to 1) calculate the overall *in silico* confirmation rate of the SRS GATK-SV calls and 2) validate the Parkinson’s Disease associated SRS GATK-SV of interest, Truvari^41^ v3.1.2 was run using the default parameters along with the --pctsim=0 parameter to turn off sequence comparison. To note, Truvari only classifies SVs as “confirmed” if the SV type (e.g., INS, DEL, DUP) is an exact match between the two callsets being compared. Because the naming of SVs was different between the two tools (i.e., with GATK-SV the type is named SVTYPE=DUP and with Sniffles2 SVTYPE=INS) before we ran Truvari all SVTYPE=DUPs in the GATK-SV callset were converted to SVTYPE=INS.

#### Data availability

For the PPMI cohort, the SV genotypes are available at the LONI IDA. For the NABEC cohort, the SV genotypes are available from the dbGaP along with the short-read sequencing WGS and SNV/small InDel genotypes, study accession phs002636.v1.p1. Statistical analysis pipeline is publicly available at https://github.com/neurogenetics/PD_SV_SRS.

## Results

### Short-Read Sequencing data Structural Variant discovery and distribution

Using the largest Parkinson’s Disease WGS SRS dataset available, we surveyed 7772 samples (mean coverage ~30X). For SV discovery and genotyping we used the GATK-SV pipeline which combines calls from four SV callers; Manta, Wham, MELT and cn.MOPS to capture the main classes of SVs. Using GATK-SV a total of 366,555 SVs were genotyped. To filter for high-quality variants only calls annotated with “PASS” were extracted, leaving a total call set of 227,357 SVs. In line with recent population genetic and human diseases studies that estimate that 401-10,884 SVs can be detected per SRS genome ^3,7,27,42,43^, our final call set contained on average 5,626 SVs per genome with a median of; 1361 insertions, 2991 deletions, 1194 duplications, 115 complex SVs and 11 inversions (Figure 2).

**Figure 1:**
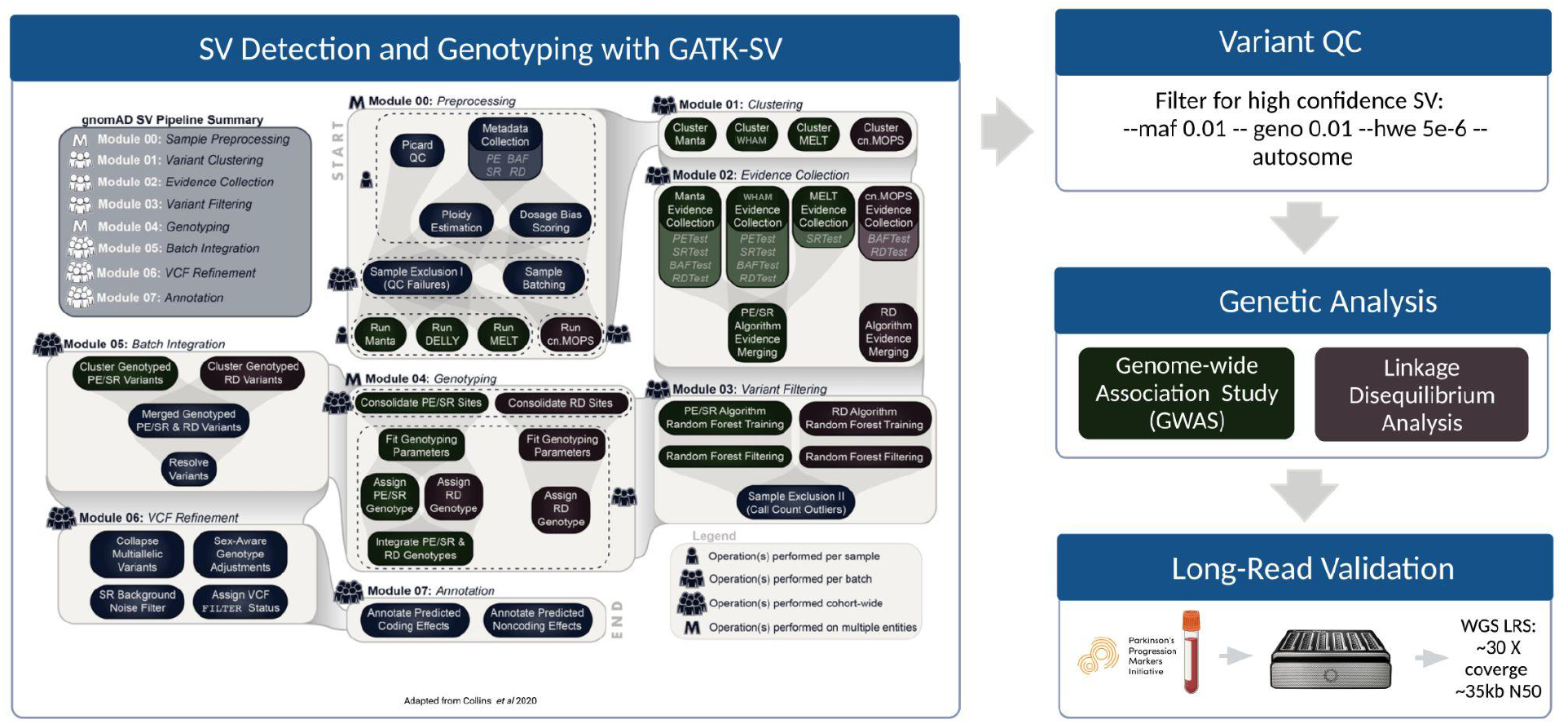
SV analysis workflow. This figure describes the study design behind the analyses included in this report.

**Figure 2:**
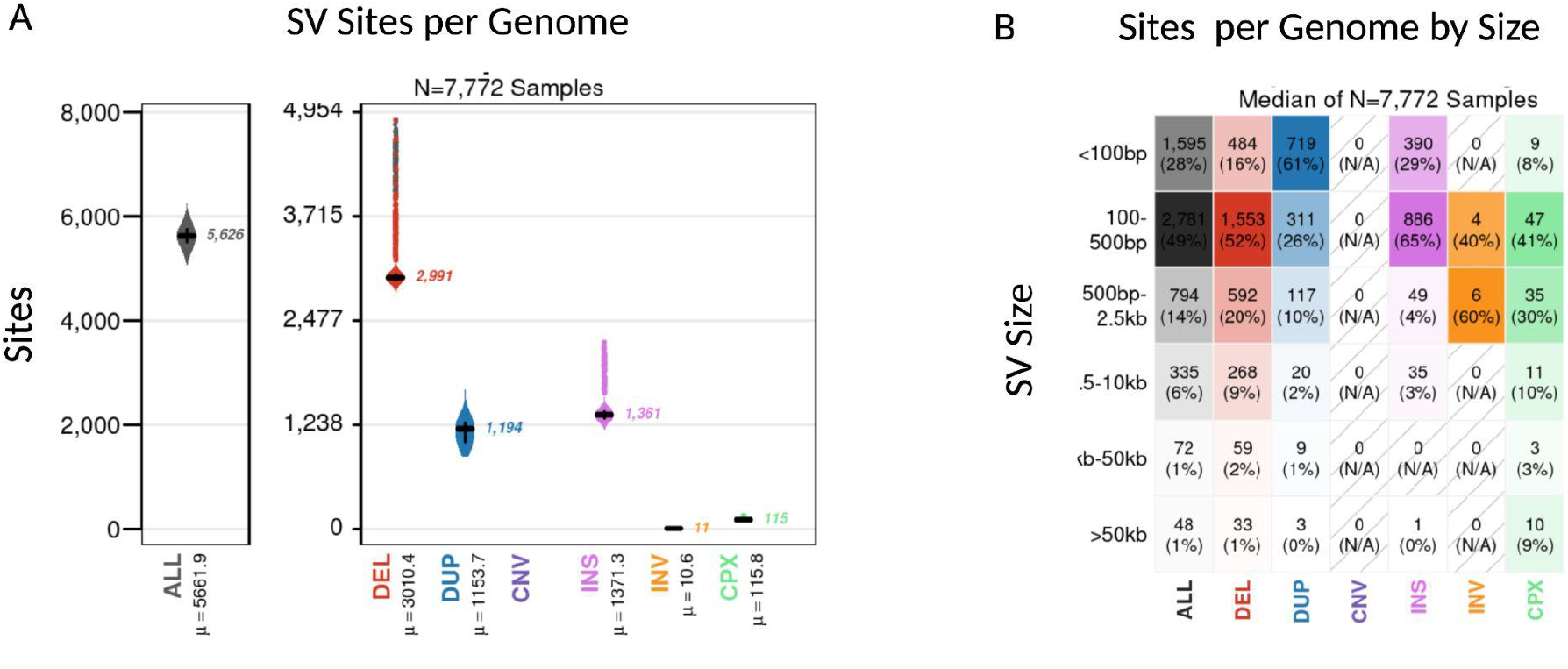
Properties of the “PASS” SVs detected in the average genome. We analyzed a total of 7772 SRS genomes after quality control. The plots show the breakdown across SV class and size. **a)** Overall on average each genome carried 5,626 SV, with a median of; 1361 insertions, 2991 deletions, 1194 duplications, 115 complex SVs and 11 inversions. **b)** The majority of SVs were small with a medium size of 329 bp. Overall only a total of 8% of SV per genome were larger than 2.5kb and 1% of SVs per genome were >50kb.

In line with previous SRS population-scale SV studies more deletions than insertions were detected in the SRS and across all SV classes the majority of the SVs were small (median 329 bp in size) with 21.40% < 100bp and 62.12% < 1kb in size (Figure 3). As expected, for the insertion size distribution we observed three main peaks at around 300bp, 2.5kb and 6kb, corresponding to *Alu*, SVA and LINE MEI^27^. The majority of the SVs were singletons (46.87%), i.e only discovered in one individual, or rare (46.69 % minor allele frequency > 0.01%).

**Figure 3:**
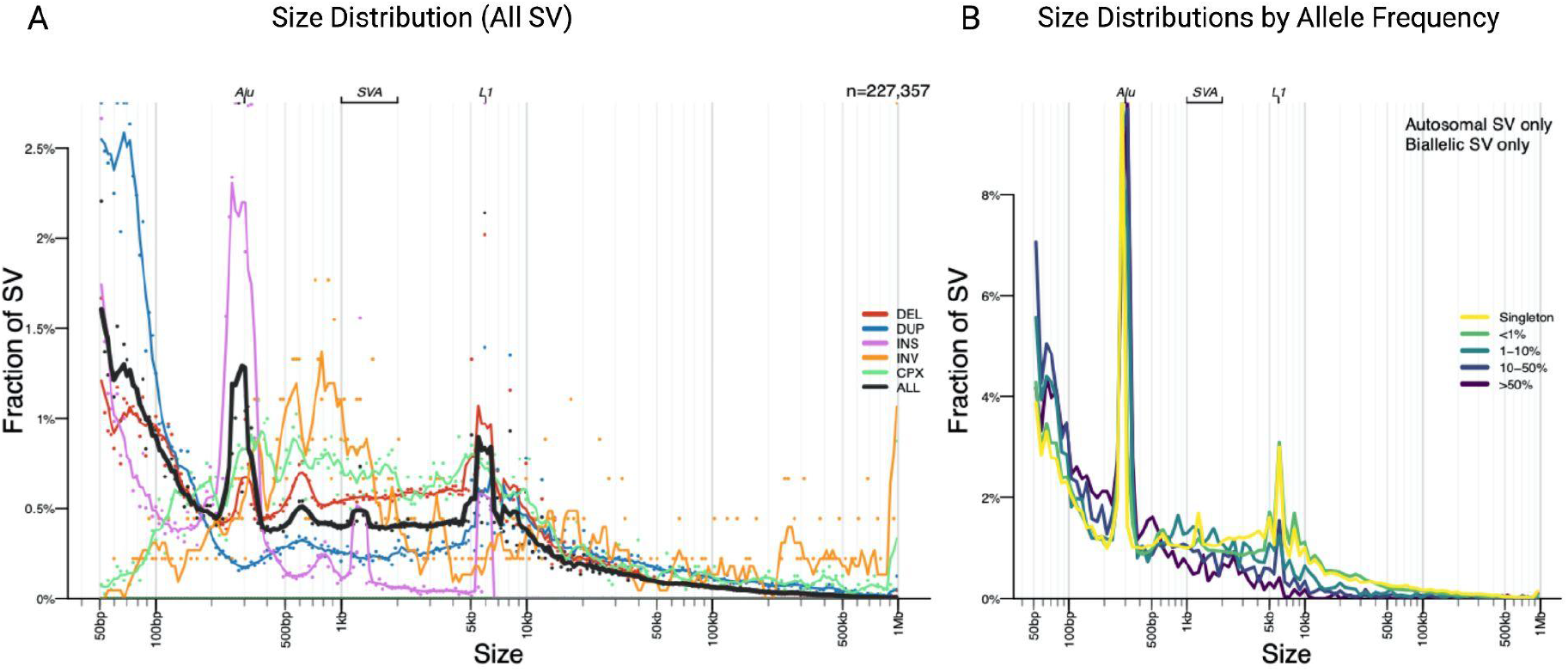
Size and allele frequency distribution of “PASS” SVs in the SRS data. **a)** The majority of SVs are small and rare. As previously reported in other large-scale SRS studies three peaks are observed at 300bp, 2kb and 6kb, representing *Alu,* SVA and LINE1 mobile element insertions respectively. b) Most SVs were singleton variants (46.87%) or rare (AF<1%)(46.69%).

### Structural Variants are candidate causal variants at known Parkinson’s Disease risk loci

The most recent Parkinson’s Disease GWAS identified 90 independent risk signals across 78 genomic regions, however, the true causal variants driving these associations are still unknown. Previous studies that integrate SV and SNV datasets have shown that SVs can tag known GWAS SNVs and are strong candidate causal variants for hundreds of human traits ^27,44,45^. In the aim of identifying SVs that may drive signals at the known Parkinson’s Disease risk loci we integrated SNVs with our GATK-SV callset. Filtering for SVs that were in linkage disequilibrium with the 90 lead Parkinson’s Disease SNVs, eight SVs were nominated.

SV detection and genotyping from SRS data can lead to many false positive calls genome-wide, especially in repetitive regions^46^. Hence it is crucial to perform validation either experimentally or with *in-silico* methods such as with matched LRS data. Here we performed extensive SV validation by generating matched LRS data for 8 PPMI individuals. We first optimized a protocol to yield high-quality LRS data from frozen human blood samples on one PromethION flow-cell. Using this protocol we generated LRS data, in which half of all sequenced base pairs (N50) belonged to reads longer than 37kb and on average had 33X coverage. SVs were detected and genotyped using Sniffles2 and the SRS and LRS SVs were compared with Truvari. A SV was considered confirmed *in-silico* if there was evidence of the SV in the corresponding LRS data and high genotype concordance across samples (defined here as > 60% of genotypes matching between SRS and LRS). Of the eight variants tested, three of the SVs were *in-silico* confirmed with the matched LRS data (Supplementary Table 2).

Two of the validated SVs; PD_DEL_chr4_14749 and PD_DEL_chr6_2338 are in moderate LD with the PD risk SNVs rs62333164 (r2=0.33, D’=0.82) and rs4140646 (r2=0.20, D’=0.65) respectively. PD_DEL_chr4_14749 is a 0.42kb intragenic deletion 35kb downstream of the gene *NEK1*. PD_DEL_chr6_2338 SV is a 0.33kb intragenic deletion 2.5kb upstream of the gene *ZSCAN9*. Both SVs are deletions of a reference *Alu* mobile element. In addition, the other validated SV PD_DEL_chr20_597 (Figure 4.A) is in strong LD with the Parkinson’s Disease risk SNV rs77351827 (r2=0.89, D’=0.95) (Figure 4.B). PD_DEL_chr20_597 is a 1.95kb intronic deletion within intron three of the gene *LRRN4* (Figure 4.C) and the deletion spans two reference *Alu* mobile elements. Further, to assess whether the SVs could drive the risk signals at these loci we attempted to run conditional analyses, however due to the current sample size of our dataset no signal existed at the three loci (Supplementary Figure 1).

**Figure 4:**
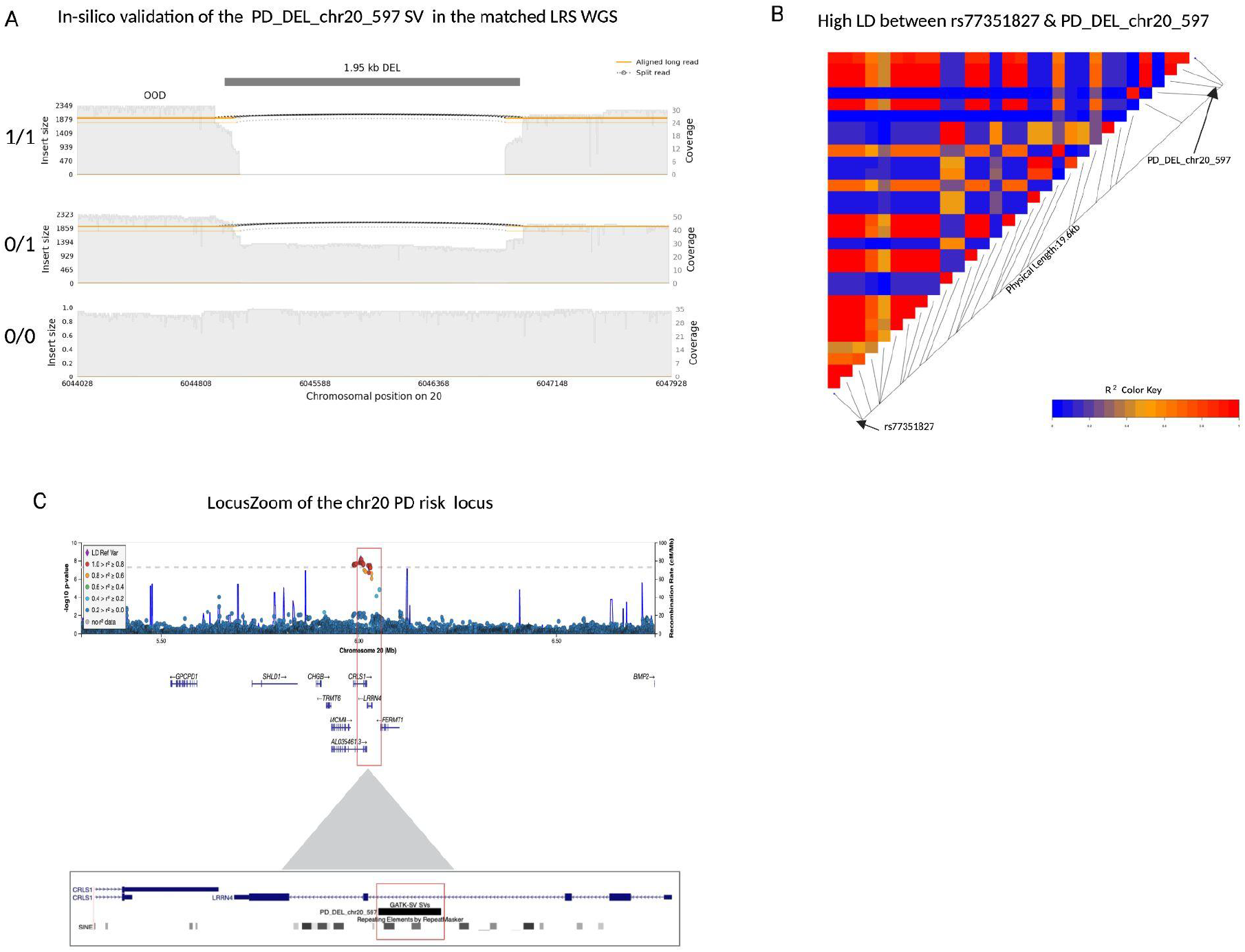
A 2kb deletion within intron 3 of *LRRN4* is a strong candidate for causal variant at the chr20 rs77351827 locus. **a)** A samplot image showing the ~2kb deletion at chr20. Aligned regions are marked in orange and the gap represents the deletion coded in black. The height of the alignment is based on the size of its largest gap. The three sequence alignment tracks follow, each alignment file plotted as a separate track in the image. The coverage for the region is shown with the gray-filled background. The SV genotypes (homozygous deletion, heterozygous deletion and homozygous reference allele/no deletion) that were predicted by GATK-SV from SRS are annotated on the left of the corresponding tracks. Each genotype was confirmed *in-silico* by the matched LRS. **b)** A LDheatmap showing pairwise LD measurements measured in R2 between the 2kb PD_DEL_chr20_597 deletion and rs77351827. High R2 values are shown in red and low R2 values in blue. PD_DEL_chr20_597 is in high LD with the lead PD risk SNV of this locus rs77351827(r2=0.89, D’=0.95). **c)** Locuszoom plot of the association signal at the chr20 rs77351827 PD risk locus from the Nalls 2019 PD SNV meta-analysis. The gene *LRRN4* lies directly under the risk signal and the schematic below shows the location of the deletion within intron 3 of the gene.

### Structural Variant Genome-wide association analysis

Despite large-scale genetic studies the majority of the common genetic variation that contributes to Parkinson’s Disease risk is still unknown. In this study we generated the first genome-wide SV dataset for Parkinson’s Disease to assess whether SVs represent part of the “missing heritability” of the disease. Following sample QC which removed samples that were; non-European, related or that carried known Parkinson’s Disease mutations and variant QC which extracted common (MAF >0.01%) autosomal biallelic SVs, 5364 samples and 3154 variants were leveraged for the association analysis. Using a GWAS approach, we identified a total of nine genome-wide significant association signals through our analysis of 2585 cases and 2779 controls (Supplementary Figure 2) The genome-wide inflation factor λ1000 was 1.004. (Supplementary Figure 3). However, *in-silico* confirmation using the matched LRS data and the tool Truvari, along with manual expectation of the matched SRS bams in IGV, indicated that the 9 “hits’’ were potentially false positive signals, either 1) there was no evidence of the SV in the matched LRS data or 2) the SV genotyping accuracy was low across the 8 tested LRS samples. To note, for the majority of loci, although a non-reference SV was present in that region, the genotype concordance was low across samples, so we could not confirm the association signal. Detailed summary statistics from the SV GWAS can be found in (Supplementary Table 4).

Of interest are two large deletions (PD_DEL_ch17_4739 and PD_DEL_chr17_4744) reaching statistical significance at the *MAPT* 17q.21.31 locus. This locus lies within a 1.5Mb inversion region of high LD conferring two distinct haplotypes H1 and the inverted H2 haplotype. The major haplotype H1 has been associated with risk of Parkinson’s Disease^47^. While these deletions do not validate in the LRS data the variants are in high LD with the H1/H2 tagging SNV rs8070723 (PD_DEL_ch17_4739; r2=0.98, D’=0.99 and PD_DEL_chr17_4744 r2=0.98, D’=0.99) suggesting that they are artifacts that likely represent the known large inversion in this region with correct genotyping accuracy but inaccurate SV detection.

Further analysis of the MAPT region identified that a 675kb large inversion variant (PD_CPX_chr17_84) is present in the non-filtered SRS GATK-SV callset, however it was removed from downstream analyses due to the Hardy Weinberg filtering. If this inversion is the known large inversion in the MAPT region it should be in complete LD with rs8070723 the H1/H2 proxy SNV. However, the LD was r2=0.65 and D’=0.90, suggesting that the genotyping accuracy of this large inversion is low in the SRS callset as well as inaccurate SV detection given the difference in size. We next aimed to investigate carriers of the inversion in the LRS data. Two of the LRS samples were predicted to carry the inversion based on rs8070723 and PD_CPX_chr17_84 genotypes. Analysis of this region in the LRS genomes identified that the two predicted carriers in fact carried a 150kb duplication as called by Sniffles2. This suggests an artifact based detection on the large inversion with inaccurate SV detection but with correct genotype like that of the two deletion SVs called from the SRS in the MAPT region. So, for this large inversion both SRS and LRS with current calling algorithms were unable to both accurately detect and genotype the SV.

### Long-read Sequencing is required to capture most Structural Variants in the genome

We sought to characterize the genome-wide distribution of SV in the LRS callsets. In line with recent LRS population studies that report ~25,000 SV per genome ^48,49^ each genome carried a mean of 27,277 SVs. Unlike the SRS callset which contains predominantly deletions, over half of the LRS SVs were insertions. Overall a medium of 14,481 insertions, 12,532 deletions, 43 duplications, 98 inversions and 123 translocations were discovered per genome.

To assess the sensitivity and specificity of the SRS SV callset we performed a detailed comparison of the SRS and matched LRS data with Truvari. It is important to note here that although recent benchmarking using 30X ONT data reported high accuracy with Sniffles2 (F-score 0.94)^38^, the following confirmation rates assume that the LRS SV calls represent the absolute ground truth, which will not be the case for 100% of the LRS SVs.

In order to benchmark our callset against the largest GATK-SV call available, we compared our confirmation rates to those from the Collins *et al* gnomad-sv study. For the comparison, in line with their approach we only focussed on “high-confidence” SVs. High-confidence SVs were defined as variants that had breakpoint-level read support (‘split-read’ evidence) and that did not span an annotated simple repeats or segmental duplications. When we focused on this restricted list the overall confirmation rate was ~80% for our calls, which is lower than the 94% reported in the Collins *et al* study. There are multiple factors that could contribute to this discrepancy. First, the final filtering step in GATK-SV (Module 07) removes variants based on the genotype quality across populations. Usually this step requires trio families to build the minGQ filtering model. However we used a model pre-trained with the 1000 Genomes samples as there were no trios present in our SRS WGS dataset. Although we implemented a very stringent final FDR cut off (1%) for our calls, as expected this likely leads to a higher false-positive rate compared to a filtering model based on family data. Another factor that likely contributes to the confirmation rate differences is that different LRS platforms, coverage and comparison tools were used. For example we used 30X ONT LRS data and ran the comparison with Truvari, compared to an analysis of 60X PacBio LRS data compared to SRS with VaPor. Therefore, given that ONT data has a higher error rate and the PacBio data has higher coverage this may explain the difference in SV discovery.

If we expand the comparison and focus on all the “PASS” SRS SVs that were used in the downstream genetic analysis reported in this present study, on average 72% of the tested SVs were confirmed in the LRS (Figure 5). A SV was classed as “confirmed” if there was evidence of that variant in the matched LRS data. For the confirmed SVs, on average the genotype concordance was 78% per genome (i.e. 78% of the genotypes matched in the SRS and LRS for those variants). As expected the majority of confirmed SVs were deletions (85% of the tested SRS deletions were confirmed). In line with the recent studies that highlight the difficulty of calling SV in repetitive regions with SRS^46^, as expected duplications drove the false positive rate, representing 40% of the false positives in the SRS callset. When we assessed the overlap between the LRS and SRS, 84% of the LRS SV were not present in the SRS callset and the majority (58%) of the SV detected solely by LRS were insertions (Figure 5).

**Figure 5:**
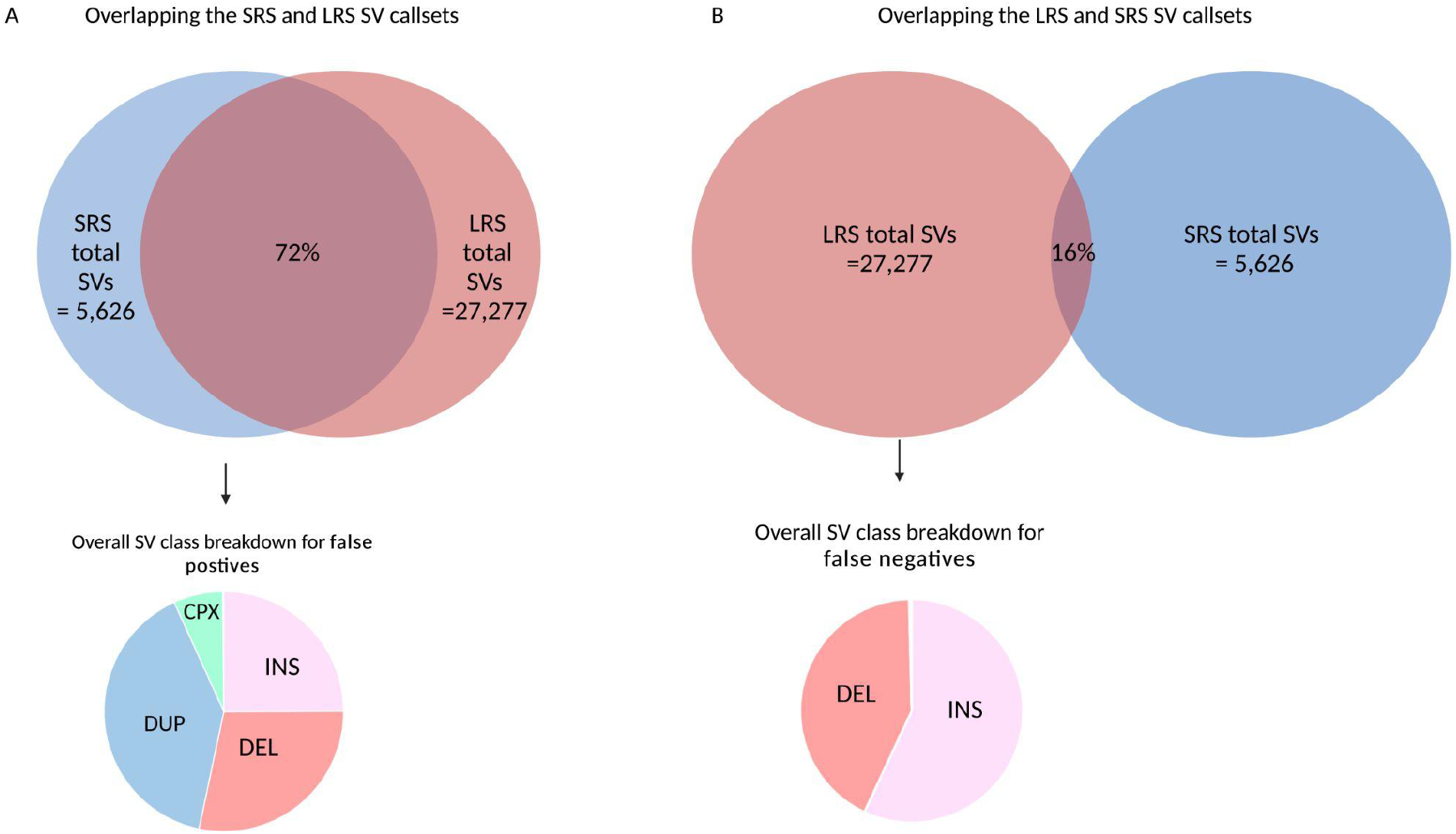
Comparison of SVs called with SRS and LRS in the eight matched PPMI blood samples. ONT LRS detects significantly more SV than SRS on average per genome. **a)** On average 5,626 SVs were detected per SRS genome compared to 27,277 with LRS. Of the 5,626 SRS SVs, 72% of the SV were confirmed *in-silico* with LRS. As expected, duplications drove the false positive rate. **b)** The majority of the SV in the genome cannot be detected with SRS alone. Of the 27,277 SVs detected with LRS, only 14% of the SVs were present in the SRS callset. The majority of these false negative calls, i.e SVs that were detected by LRS but not present in the SRS callset were insertions.

### Rare variants within reported Parkinson’s Disease causal or high risk genes

Although the primary focus of this present study was to characterize the role of common variants in Parkinson’s Disease given the utility of this new SV dataset, we aimed to report rare SV of potential interest. To date, rare variants in more than 20 genes have been reported to cause Parkinson’s Disease^50^. We explored this in our SV callset by extracting SVs within these genes that were only present in Parkinson’s Disease cases. A total of 106 rare variants lay within these genes in cases only (Supplementary Table 5). It’s important to stress here that this list of SVs were not validated and based on the rate of validation between our SRS and LRS SV callset some of the variants may be false positives.

## Discussion

Gaining a full understanding of the genetic architecture underpinning Parkinson’s Disease is critical for the development and application of therapeutic treatments that could slow or stop disease. Although progress has been made in identifying genetic factors associated with Parkinson’s Disease risk, most of the common variants driving disease have yet to be identified and even for well-characterized loci, the identity of the functional effector variant remains unknown. One reason for this is that previous genetic studies have focused solely on SNVs which represent only a fraction of the genetic variation in the human genome. SVs are a major source of genetic diversity but assessing their role in disease has been a challenge with existing genomic technologies, this type of variability, while widespread and of significant functional consequence, has been difficult to assay accurately, even with the evolution of short read whole genome sequencing^51^. Here we used GATK-SV, a multi-algorithm SV discovery pipeline for SRS WGS and report the first comprehensive genome-wide analysis of SVs in Parkinson’s Disease to date. We characterized “high confidence” common SVs in 7772 SRS genomes, representing over 412 million nucleotides of non-reference genetic variation and validated three SVs associated with Parkinson’s Disease risk at known Parkinson’s Disease risk loci.

A major bottleneck in genetics is determining the true causal variant(s) and functionally affected gene(s) within the associated risk loci. Consequently, only a few Parkinson’s Disease risk loci have been functionally validated and these mainly consist of genes that are known to cause monogenic forms of Parkinson’s Disease. One of the main motivations of this present study was to integrate SV data at these loci with the hope that by having a more complete understanding of the genetic variation at these regions we would gain a more complete understanding of genetic variation at risk loci, in turn providing insight into the biology driving risk of disease. Although this initial work is the largest and most complete assessment of SV in Parkinson’s Disease to date, it is still of relatively modest size for genetic discovery. However, we identified three SVs that are located at a known risk loci in strong LD with the lead risk SNV at each locus. Of interest, all three variants were deletions of one or more reference *Alu* mobile elements. *Alu* mobile elements are usually ~ 300 bp in length and constitute ~11% of the human reference genome with over one million copies^52^. Recent studies have shown that presence/absence variation within reference *Alu* elements can have a profound functional impact ^53,54^.

This study suggests that a 2kb deletion within intron 3 of the gene *LRRN4* (Leucine-rich repeat neuronal protein-4) is a strong candidate for causal variant at the Parkinson’s Disease risk locus at chromosome 20 (Locus 77 -https://pdgenetics.shinyapps.io/GWASBrowser/). LRRN4 is a type I transmembrane protein that is a member of the LRRN family. Previous studies report that it is expressed in the lung, heart, ovary, hippocampus and cortex and suggest it may be involved in hippocampus dependent memory retention in mice^55^. Clearly, it will be important to further understand the importance of these and other yet to be identified SV’s on risk loci, and this will likely require a combination of higher powered genetic investigation, along with the integration of functional modalities to include genomic and transcriptomic regulatory assays.

Although this study marks a significant step forward for cataloging structural variation in Parkinson’s Disease, indicative of any SV analysis from SRS data, it has several limitations. Firstly, very stringent variant QC parameters were used in an attempt to reduce the false positive rate of the SRS SV calls. Therefore it is possible that disease relevant SVs detectable via SRS may have been filtered out from the downstream analyses. Second, as shown by our LRS and SRS comparison, most SVs are not detectable using SRS alone. Meaning in the present study we have only been able to assess a small fraction of the SVs present in each genome. Taken together these factors highlight that this study likely represents a massive underestimate of the contribution of SV to risk of Parkinson’s Disease.

Generating population-scale LRS datasets to capture SVs that are currently hidden from traditional methods is an essential step towards solving the architecture of complex genetic disorders^56^. For neurodegenerative diseases specifically there are two large-scale initiatives underway to generate such datasets. The first is a Global Parkinson’s Genetics Program (GP2 https://www.gp2.org) led initiative. GP2 is the first supported resource project of the Aligning Science Across Parkinson’s, an audacious effort supporting Parkinson’s Disease research ^57^. Through this endeavor, GP2 will LRS ~1000 Parkinson’s Disease cases and control blood samples. The second initiative is led by the NIH Center of Alzheimer’s Dementias and Related Dementias (CARD https://card.nih.gov), whereby CARD is generating LRS in a total of ~ 4000 brain samples to catalog SVs in Alzheimer’s disease, Lewy Body Dementia, Frontotemporal Dementia and neurologically healthy controls.

## Supporting information

https://docs.google.com/document/d/1lQLZL8VaKCHawU8TTwTqjAsFCumk4cdjhhtOlSyT_I8/edit?usp=sharing

## Abbreviations

PD: Parkinson’s disease
SVs: Structural variants
SNVs: Single nucleotide variants
ONT: Oxford Nanopore Technologies
SRS: Short-read sequencing
LRS: Long-read sequencing
GWAS: Genome-wide association study
DEL: Deletions
DUP: Duplications
INS: Insertions
INV: Inversions
CTX: Translocations
CPX: Complex structural variants
CNV: Copy number variants
MEI: Mobile element insertions
SVA: SINE-VNTR-*Alu*
HBS: Harvard Biomarkers Study
NABEC: North American Brain Expression Consortium
PDBP: Parkinson’s Disease Biomarker Program
PPMI: Parkinson’s Progression Markers Initiative
UKBEC: United Kingdom Brain Expression Consortium
WGS: Whole genome sequencing

## Acknowledgements

We would like to thank all of the participants who donated their time and biological samples to be a part of this study. This work was supported in part by the Intramural Research Programs of the National Institute on Aging (NIA) and the National Institute of Neurological Disorders and Stroke (NINDS), part of the National Institutes of Health, Department of Health and Human Services; project numbers Z01-AG000949, 1ZIANS003154.

**AMP-PD:** Data used in the preparation of this article were partly obtained from the Accelerating Medicine Partnership® (AMP®) Parkinson’s Disease (AMP PD) Knowledge Platform. For up-to-date information on the study, visit https://www.amp-pd.org. The AMP® PD program is a public-private partnership managed by the Foundation for the National Institutes of Health and funded by the National Institute of Neurological Disorders and Stroke (NINDS) in partnership with the Aligning Science Across Parkinson’s (ASAP) initiative; Celgene Corporation, a subsidiary of Bristol-Myers Squibb Company; GlaxoSmithKline plc (GSK); The Michael J. Fox Foundation for Parkinson’s Research; Pfizer Inc.; Sanofi US Services Inc.; and Verily Life Sciences. ACCELERATING MEDICINES PARTNERSHIP and AMP are registered service marks of the U.S. Department of Health and Human Services. Clinical data and biosamples used in preparation of this article were obtained from the Michael J. Fox Foundation (MJFF) and National Institutes of Neurological Disorders and Stroke (NINDS) BioFIND study, Harvard Biomarkers Study (HBS), the NINDS Parkinson’s disease Biomarkers Program (PDBP) and MJFF Parkinson’s Progression Marker Initiative (PPMI). BioFIND is sponsored by The Michael J. Fox Foundation for Parkinson’s Research (MJFF) with support from the National Institute for Neurological Disorders and Stroke (NINDS). The BioFIND Investigators have not participated in reviewing the data analysis or content of the manuscript. For up-to-date information on the study, visit michaeljfox.org/news/biofind. The Harvard NeuroDiscovery Biomarker Study (HBS) is a collaboration of HBS investigators [full list of HBS investigator found at https://www.bwhparkinsoncenter.org/biobank/ and funded through philanthropy and NIH and Non-NIH funding sources. The HBS Investigators have not participated in reviewing the data analysis or content of the manuscript. PPMI – a public-private partnership – is funded by the Michael J. Fox Foundation for Parkinson’s Research and funding partners, a full list of the PPMI funding partners can be found at https://www.ppmi-info.org/fundingpartners. The PPMI Investigators have not participated in reviewing the data analysis or content of the manuscript. For up-to-date information on the study, visit https://www.ppmi-info.org. Parkinson’s Disease Biomarker Program (PDBP) consortium is supported by the National Institute of Neurological Disorders and Stroke (NINDS) at the National Institutes of Health. A full list of PDBP investigators can be found at https://pdbp.ninds.nih.gov/policy. The PDBP Investigators have not participated in reviewing the data analysis or content of the manuscript. **Wellderly**: This work is supported by Scripps Research Translational Institute, an NIH-NCATS Clinical and Translational Science Award (CTSA; 5 UL1TR002550). **UKBEC**: Consortium members include; Juan A. Botía, University of Murcia & UCL Great Ormond Street Institute of Child Health, Karishma D’Sa, Crick Institute, Paola Forabosco, Istituto di Ricerca Genetica e Biomedica, Italy, Sebastian Guelfi, Verge Genomics & UCL Great Ormond Street Institute of Child Health, Adaikalavan Ramasamy, Singapore Institute for Clinical Sciences, Regina H. Reynolds, UCL Great Ormond Street Institute of Child Health, Colin Smith, The University of Edinburgh, Daniah Trabzuni, UCL Queen Square Institute of Neurology, Robert Walker, The University of Edinburgh, Michael E. Weale, Genomics Plc, Oxford UK. This work was supported by the UK Dementia Research Institute which receives its funding from DRI Ltd, funded by the UK Medical Research Council, Alzheimer’s Society and Alzheimer’s Research UK.Medical Research Council (award number MR/N026004/1) and Medical Research Council (award number MR/N026004/1). **BLSA**: We would like to thank the Baltimore Longitudinal Study of Aging (BLSA) for providing control genomes and clinical data (https://www.blsa.nih.gov).The BLSA is currently supported by the Intramural Research Program of the NIH, National Institute on Aging. **LNG Path confirmed**: We are grateful to the Banner Sun Health Research Institute Brain and Body Donation Program of Sun City, Arizona for the provision of human biological materials (or specific description, e.g. brain tissue, cerebrospinal fluid). The Brain and Body Donation Program has been supported by the National Institute of Neurological Disorders and Stroke (U24 NS072026 National Brain and Tissue Resource for Parkinson’s Disease and Related Disorders), the National Institute on Aging (P30 AG19610 Arizona Alzheimer’s Disease Core Center), the Arizona Department of Health Services (contract 211002, Arizona Alzheimer’s Research Center), the Arizona Biomedical Research Commission (contracts 4001, 0011, 05-901 and 1001 to the Arizona Parkinson’s Disease Consortium) and the Michael J. Fox Foundation for Parkinson’s Research. We thank the NIH NeuroBioBank for the provision of tissue samples. **NABEC**: We thank members of the North American Brain Expression Consortium (NABEC) for providing samples derived from brain tissue. Brain tissue for the NABEC cohort were obtained from the Baltimore Longitudinal Study on Aging at the Johns Hopkins School of Medicine, the NICHD Brain and Tissue Bank for Developmental Disorders at the University of Maryland, the Banner Sun Health Research Institute Brain and Body Donation Program, and from the University of Kentucky Alzheimer’s Disease Center Brain Bank. This research was supported, in part, by the Intramural Research Program of the National Institutes of Health (National Institute on Aging, National Institute of Neurological Disorders and Stroke; project numbers 1ZIA‐NS003154, Z01‐AG000949‐02, Z01‐ES101986, and UK ADC NIA P30 AG072946.

