## Supplementary material for "Genome-wide analysis of Structural Variants in Parkinson’s Disease using Short-Read Sequencing data": https://docs.google.com/document/d/1lQLZL8VaKCHawU8TTwTqjAsFCumk4cdjhhtOlSyT_I8/edit?usp=sharing

### Supplementary Materials

#### Supplementary Figures

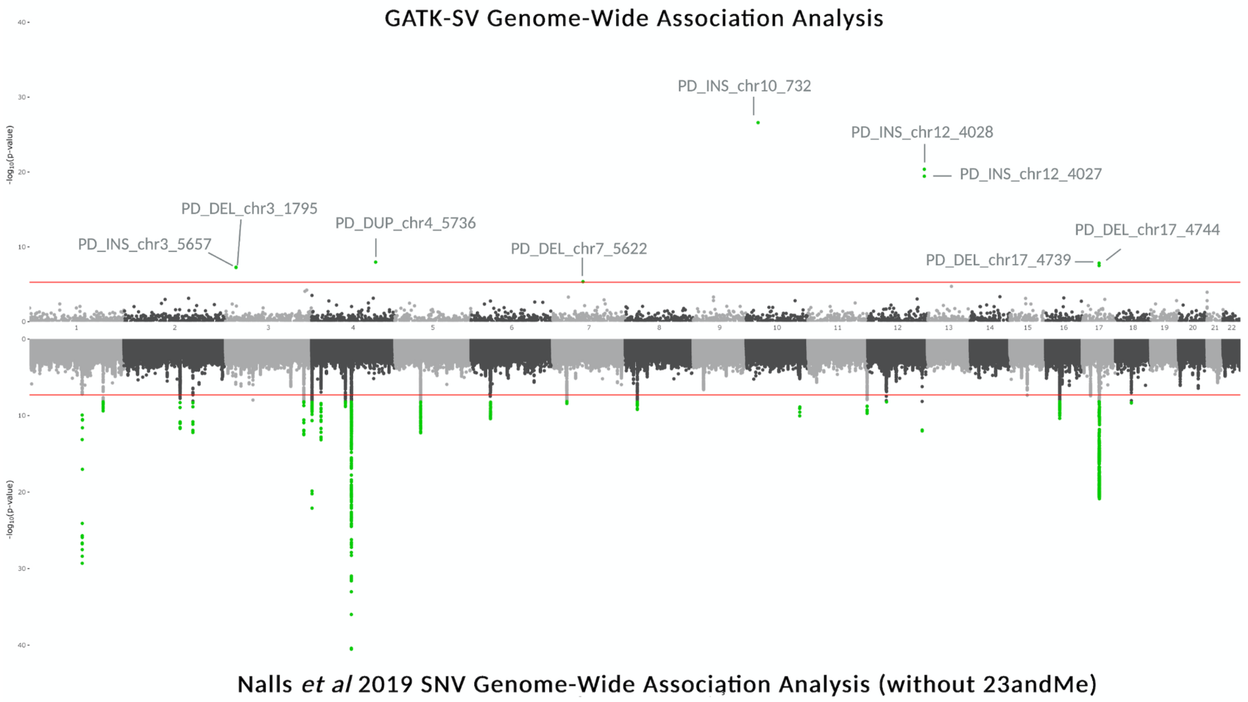

**Sup Fig 1: Manhattan (on the hudson) plots for the SV GWAS (top) and Nalls *et al* 2019 SNV (bottom) GWAS:** The top plot represents the SV GWAS results in this present study showing nine genome-wide significant SVs. Significant SVs are highlighted in green and annotated by SV ID in gray. The red line represents the genome-wide significance level ($P = 5 \times 10{-}6$) for SV. To note, none of the significant SV were confirmed with LRS. The bottom plot represents the SNV Meta5 PD GWAS. The red dotted line represents the genome-wide significance level ($P = 5 \times 10{-}8$) for SNV. The X axis represents the base pair position of variants from smallest to largest per chromosome (1–22).

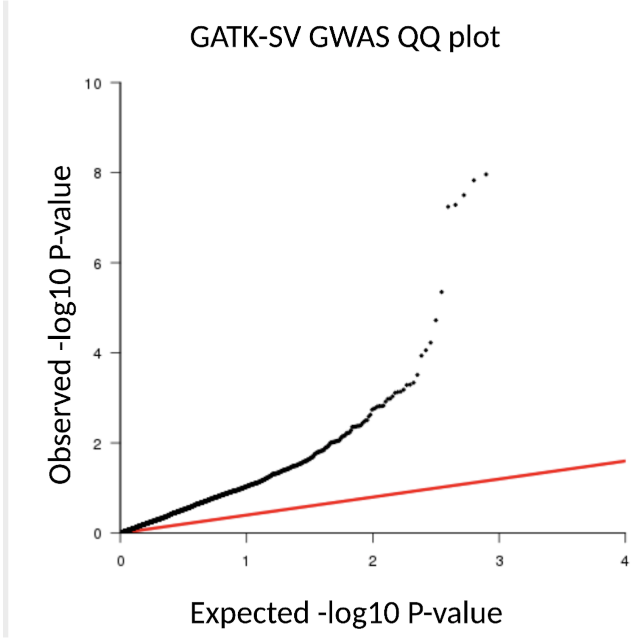

**Sup Fig 2: QQ-plot of the SV GWAS, testing 3154 variants, lambda = 1.011**

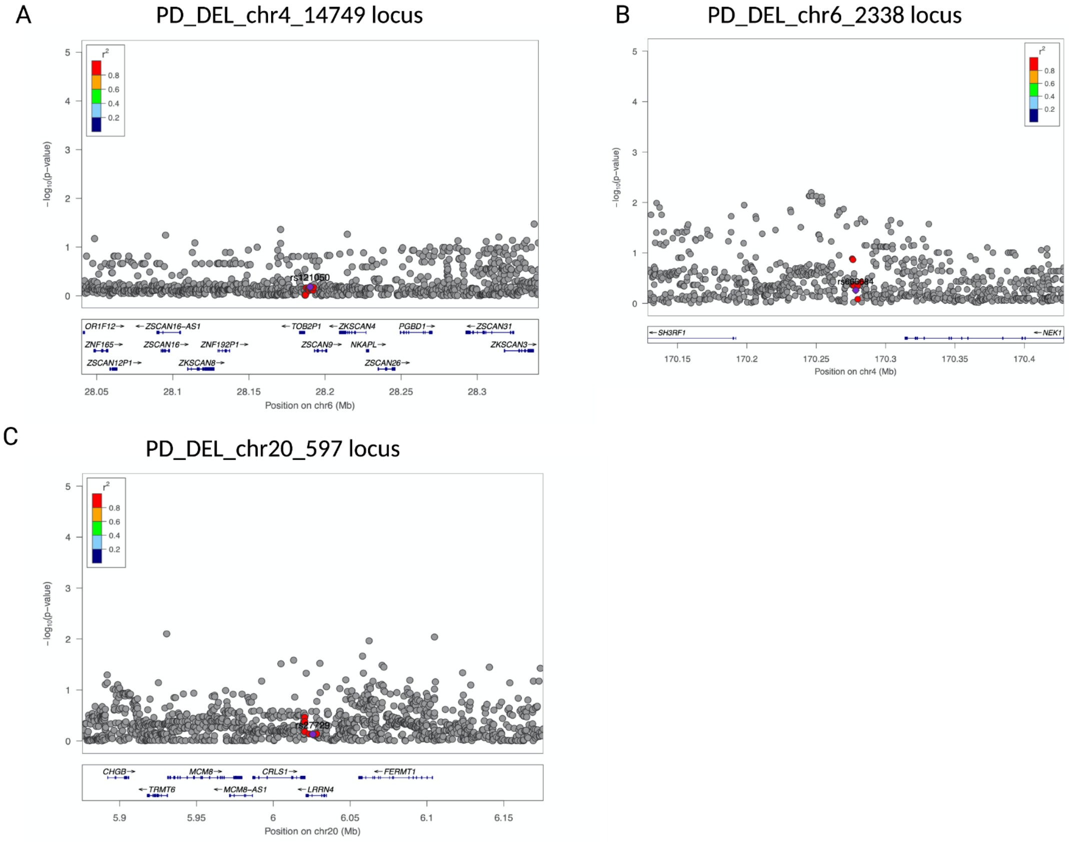

**Sup Fig 3: The risk association signal at the three known risk that contain a validated SV in linkage disequilibrium with the lead PD risk SNV** a) Locus zoom plot of the chr 4 rs62333164 locus b) Locus zoom plot of the chr6 rs4140646 locus c) Locus zoom plot of the chr20 rs77351827 locus

### Supplementary Tables

#### Supplementary Table 1: Clinical and demographic characteristics of WGS data for samples included in the SV GWAS

| Study | Cases (N) | Controls (N) | Total (N) | Female case (%) | Female control (%) | Case age at onset (mean , SD in years) | Control age at last exam (mean, SD in years) |
|---|---|---|---|---|---|---|---|
| **PPMI** | 395 | 176 | 571 | 33.92 | 11.03 | 61.56 (9.47) | 68.47 (10.21) |
| **PDBP cases & controls** | 776 | 391 | 1167 | 35.82 | 18.25 | 59.79 (10.03) | 63.49 (10.68) |
| **HBS cases & controls** | 618 | NA | 618 | 35.44 | NA | 62.04 (10.47) | NA |
| **BioFind cases & non-cases** | 95 | 68 | 163 | 34.74 | 18.40 | 61.42 (6.63) | 69.88 (7.34) |
| **NABEC controls** | NA | 253 | 253 | NA | 38.34 | NA | 54.38 (24.48) |
| **UKBEC controls** | NA | 72 | 72 | NA | 23.61 | NA | 56.88 (18.63) |
| **PD Clinical cases** | 497 | NA | 497 | 39.84 | NA | 55.71 (11.61) | NA |
| **PD Path cases** | 204 | NA | 204 | 36.76 | NA | 78.77 (8.72) | NA |
| **Wellderly controls** | NA | 1104 | 1104 | NA | 59.96 | NA | 85.03 (4.28) |
| **BLSA controls** | NA | 715 | 715 | NA | 47.55 | NA | 76.51 (11.46) |
| TOTAL | **2585** | **2779** | **5364** | | | | |
| SD: standard deviation; NA: Not available | | | | | | | |

**Supplementary Table 2: Summary of the validated SV in LD with PD risk variants**

| GATK-SV_ID | TYPE | LENGTH | NEAREST GENE | CHR | START | STOP | PD RISK SNV TAG LD | R2 | D' | DISTANCE SV TO SNV bp | LRS VALIDATED |
|---|---|---|---|---|---|---|---|---|---|---|---|
| PD_DEL_chr4_14749 | DEL | 417 | NEK1 | chr4 | 169357313 | 169357730 | rs62333164 | 0.331166 | 0.81627 | 304,276 | Yes |
| PD_DEL_chr6_2338 | DEL | 325 | ZSCAN9 | chr6 | 28222429 | 28222754 | rs4140646 | 0.203208 | 0.654587 | 451,407 | Yes |
| PD_DEL_chr20_597 | DEL | 1950 | LRRN4 | chr20 | 6045003 | 6046953 | rs77351827 | 0.889976 | 0.952148 | 21,558 | Yes |

#### Supplementary Table 3: LRS ONT Sequencing run information for the PPMI samples

| | PPMI Blood LRS | | | | |
|---|---|---|---|---|---|
| **Sample ID** | **n50** | **Total output GB** | **Chimera Rate%** | **Total Coverage** | **"PASS" SV discovered with Sniffles2 in LRS** |
| **PPMI3404** | 34.27 | 118.74 | 1.69 | 30.33 | 33362 |
| **PPMI4018** | 34.81 | 107.87 | 1.68 | 27.43 | 37506 |
| **PPMI3223** | 35.31 | 122.74 | 1.34 | 32.20 | 33084 |
| **PPMI3173** | 33.43 | 148.27 | 1.94 | 38.47 | 33159 |
| **PPMI3150** | 39.06 | 127.36 | 2.00 | 34.76 | 32783 |
| **PPMI4011** | 42.62 | 111.95 | 1.78 | 32.58 | 33260 |
| **PPMI3951** | 41.05 | 104.37 | 3.04 | 30.69 | 32901 |
| **PPMI3469** | 34.31 | 130.48 | 2.02 | 38.16 | 33027 |
| **Average=** | 36.86 | 121.47 | 1.94 | 33.08 | 33635 |

#### Supplementary Table 4: Summary statistics from the SV GWAS

Due to formatting and display issues for large tables, full summary statistics for the GATK-SV can be found here https://docs.google.com/spreadsheets/d/1aF0FDtfDWevTBMxlETK4Y2vCh4mhn0vf6C0kQ3o0TAs/edit?usp=sharing .

#### Supplementary Table 5: Rare variants only present in cases within reported PD causal or high risk genes (not validated)

| Chr | Start | Stop | SV ID | Length | Type | PD gene | Allele count in cases (0 alleles in controls) | Validated? |
|---|---|---|---|---|---|---|---|---|
| chr6 | 162075963 | 162075964 | PD_INS_chr6_5262 | 5473 | INS | PRKN | 5 | no |
| chr6 | 162450242 | 162450414 | PD_DUP_chr6_7070 | 172 | DUP | PRKN | 4 | no |
| chr15 | 61862737 | 61868437 | PD_DEL_chr15_3519 | 5700 | DEL | VPS13C | 3 | no |
| chr6 | 161717179 | 161721494 | PD_DEL_chr6_13750 | 4315 | DEL | PRKN | 3 | no |
| chr6 | 161747555 | 162202352 | PD_DEL_chr6_13758 | 454797 | DEL | PRKN | 3 | no |
| chr6 | 162090733 | 162375645 | PD_DEL_chr6_13807 | 284912 | DEL | PRKN | 3 | no |
| chr6 | 162618986 | 162629330 | PD_DUP_chr6_7080 | 10344 | DUP | PRKN | 3 | no |
| chr6 | 162075963 | 162076014 | PD_INS_chr6_5261 | 5473 | INS | PRKN | 3 | no |
| chr6 | 162075334 | 162085920 | PD_DEL_chr6_13805 | 10586 | DEL | PRKN | 2 | no |
| chr6 | 162144897 | 162144962 | PD_DEL_chr6_13814 | 65 | DEL | PRKN | 2 | no |
| chr6 | 162219198 | 162219318 | PD_DEL_chr6_13833 | 120 | DEL | PRKN | 2 | no |
| chr6 | 162226958 | 162426701 | PD_DEL_chr6_13837 | 199743 | DEL | PRKN | 2 | no |
| chr6 | 162585812 | 162588662 | PD_DEL_chr6_13908 | 2850 | DEL | PRKN | 2 | no |
| chr1 | 65380647 | 65380698 | PD_INS_chr1_1860 | 280 | INS | DNAJC6 | 2 | no |
| chr15 | 62029798 | 62029849 | PD_INS_chr15_1297 | 280 | INS | VPS13C | 2 | no |
| chr6 | 161796397 | 161796448 | PD_INS_chr6_5245 | 280 | INS | PRKN | 2 | no |
| chr6 | 162382916 | 162382935 | PD_INS_chr6_5280 | 77 | INS | PRKN | 2 | no |

| | | | | | | | | |
|---|---|---|---|---|---|---|---|---|
| chr1 | 153476402 | 155509614 | PD_CPX_chr1_207 | 2033212 | CPX | GBA | 1 | no |
| chr21 | 23636156 | 39413570 | PD_CPX_chr21_23 | 15777414 | CPX | SYNJ1 | 1 | no |
| chr1 | 65259807 | 65279008 | PD_DEL_chr1_6606 | 19201 | DEL | DNAJC6 | 1 | no |
| chr12 | 40280778 | 40281024 | PD_DEL_chr12_3482 | 246 | DEL | LRRK2 | 1 | no |
| chr12 | 40305278 | 40305597 | PD_DEL_chr12_3487 | 319 | DEL | LRRK2 | 1 | no |
| chr15 | 61962874 | 61969000 | PD_DEL_chr15_3526 | 6126 | DEL | VPS13C | 1 | no |
| chr2 | 232711618 | 232717618 | PD_DEL_chr2_20147 | 6000 | DEL | GIGYF2 | 1 | no |
| chr2 | 232724102 | 232724583 | PD_DEL_chr2_20149 | 481 | DEL | GIGYF2 | 1 | no |
| chr21 | 32698139 | 32699175 | PD_DEL_chr21_3222 | 1036 | DEL | SYNJ1 | 1 | no |
| chr21 | 32701662 | 32701826 | PD_DEL_chr21_3223 | 164 | DEL | SYNJ1 | 1 | no |
| chr22 | 38133155 | 38134956 | PD_DEL_chr22_2526 | 1801 | DEL | PLA2G6 | 1 | no |
| chr22 | 38150534 | 38150818 | PD_DEL_chr22_2528 | 284 | DEL | PLA2G6 | 1 | no |
| chr4 | 89734140 | 89734227 | PD_DEL_chr4_8207 | 87 | DEL | SNCA | 1 | no |
| chr6 | 161552826 | 161555858 | PD_DEL_chr6_13729 | 3032 | DEL | PRKN | 1 | no |
| chr6 | 161568790 | 161574790 | PD_DEL_chr6_13732 | 6000 | DEL | PRKN | 1 | no |
| chr6 | 161645168 | 161645222 | PD_DEL_chr6_13737 | 54 | DEL | PRKN | 1 | no |
| chr6 | 161748423 | 162084070 | PD_DEL_chr6_13759 | 335647 | DEL | PRKN | 1 | no |
| chr6 | 161796690 | 161797277 | PD_DEL_chr6_13760 | 587 | DEL | PRKN | 1 | no |
| chr6 | 161856485 | 161856848 | PD_DEL_chr6_13761 | 363 | DEL | PRKN | 1 | no |
| chr6 | 161949814 | 161964160 | PD_DEL_chr6_13777 | 14346 | DEL | PRKN | 1 | no |
| chr6 | 162002000 | 162066790 | PD_DEL_chr6_13785 | 64790 | DEL | PRKN | 1 | no |
| chr6 | 162033387 | 162038806 | PD_DEL_chr6_13798 | 5419 | DEL | PRKN | 1 | no |
| chr6 | 162034265 | 162235649 | PD_DEL_chr6_13799 | 201384 | DEL | PRKN | 1 | no |
| chr6 | 162039267 | 162040666 | PD_DEL_chr6_13800 | 1399 | DEL | PRKN | 1 | no |
| chr6 | 162052790 | 162227130 | PD_DEL_chr6_13801 | 174340 | DEL | PRKN | 1 | no |
| chr6 | 162070757 | 162072119 | PD_DEL_chr6_13804 | 1362 | DEL | PRKN | 1 | no |
| chr6 | 162109710 | 162250990 | PD_DEL_chr6_13808 | 141280 | DEL | PRKN | 1 | no |
| chr6 | 162111409 | 162354299 | PD_DEL_chr6_13809 | 242890 | DEL | PRKN | 1 | no |

| | | | | | | | | |
|---|---|---|---|---|---|---|---|---|
| chr6 | 162113862 | 162115810 | PD_DEL_chr6_13810 | 1948 | DEL | PRKN | 1 | no |
| chr6 | 162121442 | 162381034 | PD_DEL_chr6_13811 | 259592 | DEL | PRKN | 1 | no |
| chr6 | 162149508 | 162164761 | PD_DEL_chr6_13815 | 15253 | DEL | PRKN | 1 | no |
| chr6 | 162167769 | 162458643 | PD_DEL_chr6_13821 | 290874 | DEL | PRKN | 1 | no |
| chr6 | 162171863 | 162315221 | PD_DEL_chr6_13823 | 143358 | DEL | PRKN | 1 | no |
| chr6 | 162174064 | 162206317 | PD_DEL_chr6_13824 | 32253 | DEL | PRKN | 1 | no |
| chr6 | 162199999 | 162278845 | PD_DEL_chr6_13830 | 78846 | DEL | PRKN | 1 | no |
| chr6 | 162215327 | 162405338 | PD_DEL_chr6_13832 | 190011 | DEL | PRKN | 1 | no |
| chr6 | 162220681 | 162380023 | PD_DEL_chr6_13835 | 159342 | DEL | PRKN | 1 | no |
| chr6 | 162227931 | 162466456 | PD_DEL_chr6_13838 | 238525 | DEL | PRKN | 1 | no |
| chr6 | 162229020 | 162229098 | PD_DEL_chr6_13839 | 78 | DEL | PRKN | 1 | no |
| chr6 | 162235983 | 162425918 | PD_DEL_chr6_13840 | 189935 | DEL | PRKN | 1 | no |
| chr6 | 162246220 | 162247864 | PD_DEL_chr6_13843 | 1644 | DEL | PRKN | 1 | no |
| chr6 | 162251287 | 162257663 | PD_DEL_chr6_13844 | 6376 | DEL | PRKN | 1 | no |
| chr6 | 162254355 | 162542940 | PD_DEL_chr6_13845 | 288585 | DEL | PRKN | 1 | no |
| chr6 | 162258574 | 162361381 | PD_DEL_chr6_13847 | 102807 | DEL | PRKN | 1 | no |
| chr6 | 162299108 | 162535581 | PD_DEL_chr6_13851 | 236473 | DEL | PRKN | 1 | no |
| chr6 | 162302274 | 162302961 | PD_DEL_chr6_13852 | 687 | DEL | PRKN | 1 | no |
| chr6 | 162313437 | 162313498 | PD_DEL_chr6_13857 | 61 | DEL | PRKN | 1 | no |
| chr6 | 162313496 | 162323202 | PD_DEL_chr6_13858 | 9706 | DEL | PRKN | 1 | no |
| chr6 | 162316450 | 162488715 | PD_DEL_chr6_13859 | 172265 | DEL | PRKN | 1 | no |
| chr6 | 162334685 | 162444403 | PD_DEL_chr6_13863 | 109718 | DEL | PRKN | 1 | no |
| chr6 | 162365311 | 162523580 | PD_DEL_chr6_13870 | 158269 | DEL | PRKN | 1 | no |
| chr6 | 162371541 | 162371652 | PD_DEL_chr6_13871 | 111 | DEL | PRKN | 1 | no |
| chr6 | 162382236 | 162384739 | PD_DEL_chr6_13875 | 2503 | DEL | PRKN | 1 | no |
| chr6 | 162440000 | 162454790 | PD_DEL_chr6_13883 | 14790 | DEL | PRKN | 1 | no |
| chr6 | 162442763 | 162442929 | PD_DEL_chr6_13884 | 166 | DEL | PRKN | 1 | no |
| chr6 | 162444717 | 162547267 | PD_DEL_chr6_13885 | 102550 | DEL | PRKN | 1 | no |
| chr6 | 162480193 | 162617780 | PD_DEL_chr6_13892 | 137587 | DEL | PRKN | 1 | no |

| | | | | | | | | |
|---|---|---|---|---|---|---|---|---|
| chr6 | 162510025 | 162514312 | PD_DEL_chr6_13899 | 4287 | DEL | PRKN | 1 | no |
| chr6 | 162670965 | 162672732 | PD_DEL_chr6_13914 | 1767 | DEL | PRKN | 1 | no |
| chr1 | 16989078 | 17061247 | PD_DUP_chr1_1640 | 72169 | DUP | ATP13A2 | 1 | no |
| chr1 | 20639135 | 20684110 | PD_DUP_chr1_1915 | 44975 | DUP | PINK1 | 1 | no |
| chr15 | 61885826 | 61885982 | PD_DUP_chr15_2618 | 156 | DUP | VPS13C | 1 | no |
| chr15 | 62009699 | 62009784 | PD_DUP_chr15_2624 | 85 | DUP | VPS13C | 1 | no |
| chr6 | 161364041 | 161364331 | PD_DUP_chr6_7023 | 290 | DUP | PRKN | 1 | no |
| chr6 | 161480293 | 161480352 | PD_DUP_chr6_7030 | 59 | DUP | PRKN | 1 | no |
| chr6 | 161567720 | 162085660 | PD_DUP_chr6_7032 | 517940 | DUP | PRKN | 1 | no |
| chr6 | 161756556 | 161756619 | PD_DUP_chr6_7038 | 63 | DUP | PRKN | 1 | no |
| chr6 | 162070917 | 162070970 | PD_DUP_chr6_7046 | 53 | DUP | PRKN | 1 | no |
| chr6 | 162117364 | 162246481 | PD_DUP_chr6_7050 | 129117 | DUP | PRKN | 1 | no |
| chr6 | 162145143 | 162147680 | PD_DUP_chr6_7051 | 2537 | DUP | PRKN | 1 | no |
| chr6 | 162461124 | 162713771 | PD_DUP_chr6_7073 | 252647 | DUP | PRKN | 1 | no |
| chr6 | 162472576 | 162503251 | PD_DUP_chr6_7074 | 30675 | DUP | PRKN | 1 | no |
| chr6 | 162634964 | 162679849 | PD_DUP_chr6_7081 | 44885 | DUP | PRKN | 1 | no |
| chr15 | 89325139 | 89325143 | PD_INS_chr15_2041 | 54 | INS | POLG | 1 | no |
| chr15 | 89325150 | 89325159 | PD_INS_chr15_2042 | 50 | INS | POLG | 1 | no |
| chr21 | 32695626 | 32695677 | PD_INS_chr21_886 | 1240 | INS | SYNJ1 | 1 | no |
| chr22 | 38197259 | 38197310 | PD_INS_chr22_656 | 55 | INS | PLA2G6 | 1 | no |
| chr3 | 132504888 | 132504939 | PD_INS_chr3_3782 | 73 | INS | DNAJC13 | 1 | no |
| chr6 | 161633443 | 161633452 | PD_INS_chr6_5237 | 281 | INS | PRKN | 1 | no |
| chr6 | 161760074 | 161760079 | PD_INS_chr6_5240 | 279 | INS | PRKN | 1 | no |
| chr6 | 161915409 | 161915460 | PD_INS_chr6_5256 | 280 | INS | PRKN | 1 | no |
| chr6 | 162008375 | 162008397 | PD_INS_chr6_5259 | 5219 | INS | PRKN | 1 | no |
| chr6 | 162195488 | 162195539 | PD_INS_chr6_5267 | 279 | INS | PRKN | 1 | no |
| chr6 | 162361022 | 162361073 | PD_INS_chr6_5279 | 280 | INS | PRKN | 1 | no |
| chr6 | 162468687 | 162468738 | PD_INS_chr6_5287 | 393 | INS | PRKN | 1 | no |
| chr6 | 162662596 | 162676889 | PD_INS_chr6_5295 | 85 | INS | PRKN | 1 | no |

| | | | | | | | | |
|---|---|---|---|---|---|---|---|---|
| chr6 | 162725077 | 162725077 | PD_INS_chr6_5298 | 192 | INS | PRKN | 1 | no |
| chr2 | 63332247 | 102880455 | PD_INV_chr2_13 | 39548208 | INV | HTRA2 | 1 | no |
| chr4 | 39626057 | 57263812 | PD_INV_chr4_9 | 17637755 | INV | UCHL1 | 1 | no |
